# The effect of Nipped-B-Like (Nipbl) haploinsufficiency on genome-wide cohesin binding and target gene expression: modeling Cornelia de Lange Syndrome

**DOI:** 10.1101/134825

**Authors:** Daniel A. Newkirk, Yen-Yun Chen, Richard Chien, Weihua Zeng, Jacob Biesinger, Ebony Flowers, Shimako Kawauchi, Rosaysela Santos, Anne L. Calof, Arthur D. Lander, Xiaohui Xie, Kyoko Yokomori

## Abstract

Cornelia de Lange Syndrome (CdLS) is a multisystem developmental disorder frequently associated with heterozygous loss-of-function mutations of *Nipped-B-like* (*NIPBL*), the human homolog of Drosophila *Nipped-B*. NIPBL loads cohesin onto chromatin. Cohesin mediates sister chromatid cohesion important for mitosis, but is also increasingly recognized as a regulator of gene expression. In CdLS patient cells and animal models, the presence of multiple gene expression changes with little or no sister chromatid cohesion defect suggests that disruption of gene regulation underlies this disorder. However, the effect of *NIPBL* haploinsufficiency on cohesin binding, and how this relates to the clinical presentation of CdLS, has not been fully investigated. Nipbl haploinsufficiency causes CdLS-like phenotype in mice. We examined genome-wide cohesin binding and its relationship to gene expression using mouse embryonic fibroblasts (MEFs) from *Nipbl* +/- mice that recapitulate the CdLS phenotype. We found a global decrease in cohesin binding, including at CCCTC-binding factor (CTCF) binding sites and repeat regions. Cohesin-bound genes were found to be enriched for histone H3 lysine 4 trimethylation (H3K4me3) at their promoters; were disproportionately downregulated in *Nipbl* mutant MEFs; and displayed evidence of reduced promoter-enhancer interaction. The results suggest that gene activation is the primary cohesin function sensitive to Nipbl reduction. Over 50% of significantly dysregulated transcripts in mutant MEFs come from cohesin target genes, including genes involved in adipogenesis that have been implicated in contributing to the CdLS phenotype. Thus, decreased cohesin binding at the gene regions directly contributes to disease-specific expression changes. Taken together, our Nipbl haploinsufficiency model allows us to analyze the dosage effect of cohesin loading on CdLS development.

## Introduction

CdLS (OMIM 122470, 300590, 610759) is a dominant genetic disorder estimated to occur in 1 in 10,000 individuals, characterized by facial dysmorphism, hirsutism, upper limb abnormalities, cognitive retardation, and growth abnormalities [1, 2]. Mutations in the *NIPBL* gene are linked to more than 55% of CdLS cases [3, 4]. NIPBL is an evolutionarily conserved, essential protein that is required for chromatin loading of cohesin [5]. Cohesin is a multiprotein complex, also conserved and essential, which functions in chromosome structural organization important for genome maintenance and gene expression [6-8]. Mutations in the cohesin subunits SMC1 (human SMC1 (hSMC1), SMC1A) and hSMC3 were also found in a minor subset of clinically milder CdLS cases (∼5% and <1%, respectively) [9-11]. More recently, mutation of HDAC8, which regulates cohesin dissociation from chromatin in mitosis, was found in a subset of CdLS patients (OMIM 300882) [12]. Mutations in the non-SMC cohesin component *Rad21* gene have also been found in patients with a CdLS-like phenotype (OMIM 606462), with much milder cognitive impairment [13]. Thus, mutations of cohesin subunits and regulators of cohesin’s chromatin association cause related phenotypes, suggesting that impairment of the cohesin pathway makes significant contributions to the disease [2, 14].

The most common cause of CdLS is *NIPBL* haploinsufficiency [2, 15, 16]. Even a 15% decrease in expression was reported to cause mild but distinct CdLS phenotype, suggesting the extreme sensitivity of human development to *NIPBL* gene dosage [17, 18]. Similarly, *Nipbl* heterozygous mutant (*Nipbl* +/-) mice display only a 25-30% decrease in *Nipbl* transcripts, presumably due to compensatory upregulation of the intact allele [19]. They, however, exhibit wide-ranging defects characteristic of the disease, including small size, craniofacial anomalies, microbrachycephaly, heart defects, hearing abnormalities, low body fat, and delayed bone maturation [19]. Thus, these results indicate a conserved high sensitivity of mammalian development to *Nipbl* gene dosage and that Nipbl +/- mice can serve as a CdLS disease model,.

Although a canonical function of cohesin is sister chromatid cohesion critical for mitosis [8], a role for cohesin in gene regulation has been argued for based on work in multiple organisms [20, 21]. The partial decrease of *Nipbl* expression in CdLS patients and *Nipbl* +/- mice was not sufficient to cause a significant sister chromatid cohesion defect or abnormal mitosis [19, 22-24]. Instead, a distinctive profile of gene expression changes was observed, revealing dosage-sensitive functional hierarchy of cohesin and strongly suggesting that transcriptional dysregulation underlies the disease phenotype [6, 18, 19, 25]. In Nipbl +/- mutant mice, gene expression changes are pervasive, though mostly minor, raising the possibility that small expression perturbations of multiple genes collectively contribute to the disease phenotype [19]. Indeed, combinatorial partial depletion of key developmental genes dysregulated in this mouse model, successfully recapitulated specific aspects of the CdLS-like phenotype in zebrafish [26]. A recent study on CdLS patient lymphoblasts and correlation with ChIP-seq revealed dysregulation of RNA processing genes, which also explains a certain aspect of CdLS cellular phenotype [27]. However, discordance of NIPBL and cohesin binding patterns in mammalian genome suggests that NIPBL may have cohesin-independent transcriptional effects [28]. Thus, it is important to determine the effects of Nipbl haploinsufficiency on cohesin binding and cohesin-bound target genes. While a similar study has been done using patient and control cells [18], the Nipbl +/- mouse model in comparison to the Nipbl +/+ wild type provides an ideal isogenic for this purpose¨

Cohesin is recruited to different genomic regions and affects gene expression in different ways in mammalian cells [6, 7, 29]. In mammalian cells, one major mechanism of cohesin-mediated gene regulation is through CTCF [30-33]. CTCF is a zinc finger DNA-binding protein and was shown to as a transcriptional activator/repressor as well as an insulator [34]. Genome-wide chromatin immunoprecipitation (ChIP) analyses revealed that a significant number of cohesin binding sites overlap with those of CTCF in human and mouse somatic cells [30, 31]. Cohesin is recruited to these sites by CTCF and mediates CTCF’s insulator function by bridging distant CTCF sites at, for example, the *H19/IGF2*, *IFNγ*, apolipoprotein, and *β-globin* loci [30, 31, 33, 35-38]. While CTCF recruits cohesin, it is cohesin that plays a primary role in long-distance chromatin interaction [36]. A more recent genome-wide Chromosome Conformation Capture Carbon Copy (5C) study revealed that CTCF/cohesin tends to mediate long-range chromatin interactions defining megabase-sized topologically associating domains (TADs) [39], indicating that CTCF and cohesin together play a fundamental role in chromatin organization in the nucleus. Cohesin also binds to other genomic regions and functions in a CTCF-independent manner in gene activation by facilitating promoter-enhancer interactions together with Mediator [35, 39-41]. Significant overlap between cohesin at non-CTCF sites and cell type-specific transcription factor binding sites was found, suggesting a role for cohesin at non-CTCF sites in cell type-specific gene regulation [41-43]. In addition, cohesin is recruited to heterochromatic repeat regions [44, 45]. To what extent these different modes of cohesin recruitment and function are affected by *NIPBL* haploinsufficiency in CdLS has not been examined.

Here, using MEFs derived from *Nipbl*+/- mice, we analyzed the effect of Nipbl haploinsufficiency on cohesin-mediated gene regulation and identified cohesin target genes that are particularly sensitive to partial reduction of Nipbl. Our results indicate that Nipbl is required for cohesin binding to both CTCF and non-CTCF sites, as well as repeat regions. Significant correlation was found between gene expression changes in Nipbl mutant cells and cohesin binding to the gene regions, in particular promoter regions, suggesting that even modest Nipbl reduction directly and significantly affects expression of cohesin-bound genes. Target genes are enriched for developmental genes, including multiple genes that regulate adipogenesis, which is impaired in Nipbl +/- mice [19]. The results indicate that Nipbl regulates a significant number of genes through cohesin. While their expression levels vary in wild type cells, the Nipbl/cohesin target genes tend on the whole to be downregulated in Nipbl mutant cells, indicating that Nipbl and cohesin are important for activation of these genes. Consistent with this, these genes are enriched for H3 lysine 4 trimethylation (H3K4me3) at the promoter regions. The long-distance interaction of the cohesin-bound promoter and a putative enhancer region is decreased by Nipbl reduction, indicating that reduced cohesin binding by Nipbl haploinsufficiency affects chromatin interactions. Collectively, the results reveal that Nipbl haploinsufficiency globally reduces cohesin binding, and its major transcriptional consequence is downregulation of cohesin target genes.

## METHODS

### Cells and antibodies

Mouse embryonic fibroblasts (MEFs) derived from E15.5 wild type and *Nipbl* mutant embryos were used as described previously [19]. In brief, mice heterozygous for *Nipbl* mutation were generated (*Nipbl* +/-) from gene-trap-inserted ES cells. This mutation resulted in a net 30-50% decrease in *Nipbl* transcripts in the mice, along with many phenotypes characteristic of human CdLS patients [19]. Wild type and mutant MEF cell lines derived from the siblings were cultured at 37°C and 5% CO_2_ in DMEM (Gibco) supplemented with 10% fetal bovine serum and penicillin-streptomycin (50U/mL). Antibodies specific for hSMC1 and Rad21 were previously described [46]. Rabbit polyclonal antibody specific for the NIPBL protein was raised against a bacterially-expressed recombinant polypeptide corresponding to the C-terminal fragment of NIPBL isoform A (NP_597677.2) (amino acids 2429–2804) [45]. Anti-histone H3 rabbit polyclonal antibody was from Abcam (ab1791).

### ChIP-sequencing (ChIP-seq) and ChIP-PCR

ChIP was carried out as described previously [35]. Approximately 50 μg DNA was used per IP. Cells were crosslinked 10 mins with 1% formaldehyde, lysed, and sonicated using the Bioruptor from Diagenode to obtain ∼200bp fragments using a 30 sec on/off cycle for 1 hr. Samples were diluted and pre-cleared for 1 hr with BSA and Protein A beads. Pre-cleared extracts were incubated with Rad21, Nipbl, and preimmune antibodies overnight. IP was performed with Protein A beads with subsequent washes. DNA was eluted off beads, reversed crosslinked for 8 hrs, and purified with the Qiagen PCR Purification Kit. Samples were submitted to Ambry Genetics (Aliso Viejo, CA) for library preparation and sequencing using the Illumina protocol and the Illumina Genome Analyzer (GA) system. The total number of reads before alignment were: preimmune IgG, 7,428,656; Rad21 in control WT, 7,200,450; Rad21 in Nipbl+/-, 4,668,622; histone H3 in WT, 26,630,000; and histone H3 in Nipbl+/-, 24,952,439. Sequences were aligned to the mouse mm9 reference genome using Bowtie (with parameters–n2, -k20, —best, —strata, —chunkmbs 384) [47]. ChIP-seq data is being submitted to GEO. PCR primers used for manual ChIP confirmation are listed in Table 1. Primers corresponding to repeat sequences (major and minor satellite, rDNA, and SINEB1 repeats) were from Matens et al. [48]. For manual ChIP-PCR analysis of selected genomic locations, ChIP signals were normalized with preimmune IgG and input DNA from each cell sample as previously described [35, 45, 49]. The experiments were repeated at least three times using MEF samples from different litters, which yielded consistent results. PCR reactions were done in duplicates or triplicates.

**Table 1.**
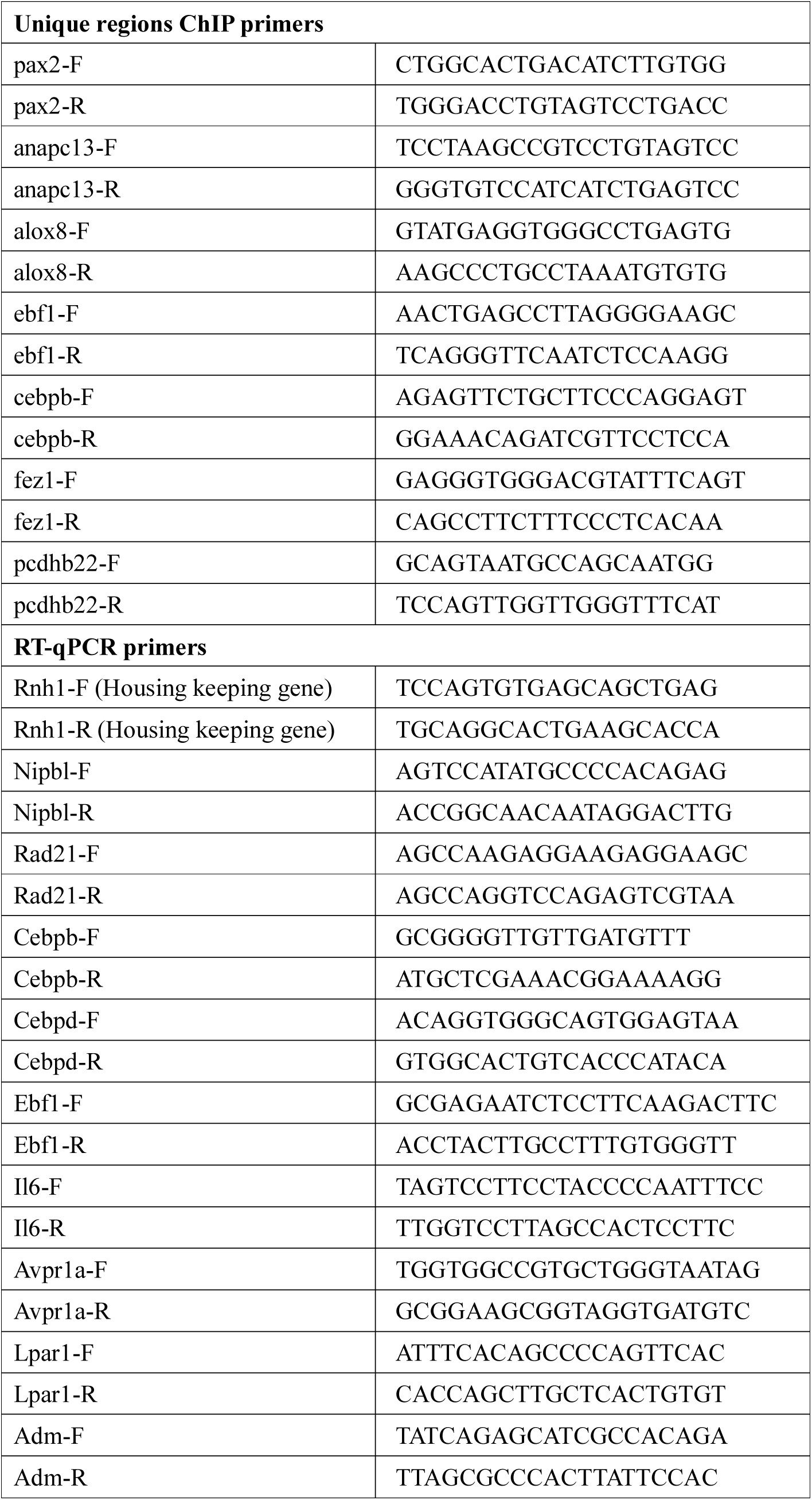
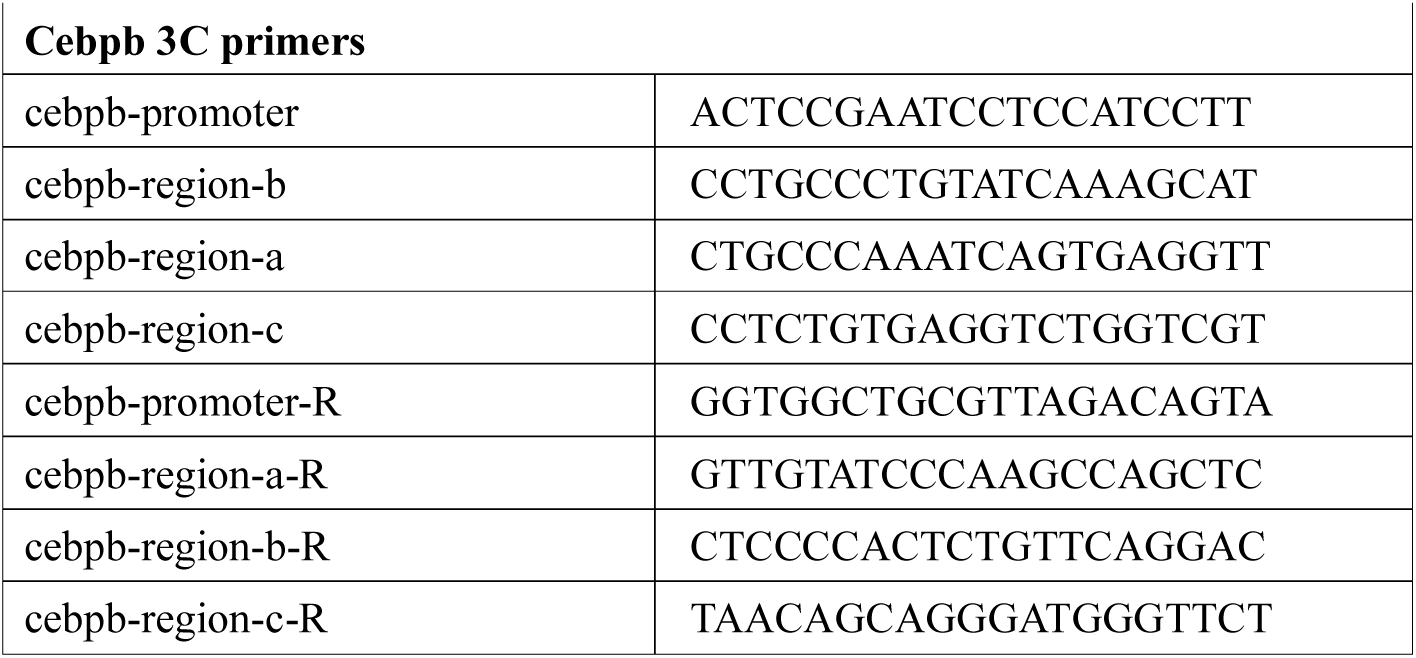
The list of PCR primers.

### Peak Finding

Peaks were called using AREM (<u>A</u>ligning ChIP-seq <u>R</u>eads using <u>E</u>xpectation <u>M</u>aximization) as previously described [50]. AREM incorporates sequences with one or many mappings to call peaks as opposed to using only uniquely mapping reads, allowing one to call peaks normally missed due to repetitive sequence. Since many peaks for Rad21 as well as CTCF can be found in repetitive sequence [50, 51], we used a mixture model to describe the data, assuming K + 1 clusters of sequences (K peaks and background). Maximum likelihood is used to estimate the locations of enrichment, with the read alignment probabilities iteratively updated using EM. Final peaks are called for each window assuming a Poisson distribution, calculating a *p*-value for each sequence cluster. The false discovery rate for all peaks was determined relative to the pre-immune sample, with EM performed independently for the pre-immune sample as well. Full algorithm details are available, including a systematic comparison to other common peak callers such as SICER and MACS [50]. Overlap between peaks and genomic regions of interest were generated using Perl and Python scripts as well as pybedtools [52, 53]. Figures were generated using the R statistical package [54]. Visualization of sequence pileup utilized the UCSC Genome Browser [55, 56].

### Motif Analysis

*De Novo* motif discovery was performed using Multiple Expectation maximization for Motif Elicitation (MEME) version 6.1 [57]. Input sequences were limited to 200 bp in length surrounding the summit of any given peak, and the number reduced to 1000 randomly sampled sequences from the set of all peak sequences. Motif searches for known motifs were performed by calculation of a log-odds ratio contrasting the position weight matrix with the background nucleotide frequency. Baseline values were determined from calculations across randomly selected regions of the genome. Randomly selected 200bp genomic regions were used to calculate a false discovery rate (FDR) at several position weight matrix (PWM) score thresholds. We chose the motif-calling score threshold corresponding to a 4.7% FDR. The *p*-values were derived for the number of matches above the z-score threshold relative to the background using a hypergeometric test.

### Expression data Analysis

Affymetrix MOE430A 2.0 array data for mouse embryonic fibroblasts (10 data sets for the wild type and nine for Nipbl+/- mutant MEFS) were previously published [19]. Expression data were filtered for probe sets with values below 300 and above 20,000, with the remainder used for downstream analysis. Differential expression and associated *p*-values were determined using Cyber-t, which uses a modified t-test statistic [58]. Multiple Hypothesis Testing correction was performed using a permutation test with 1000 permutations of the sample data. Probe sets were collapsed into genes by taking the median value across all probe sets representing a particular gene. Raw expression values for each gene are represented as a z-score, which denotes the number of standard deviations that value is away from the mean value across all genes. Gene ontology analysis was performed using PANTHER [59, 60] with a cutoff of p < 0.05.

### KS test

Genes were sorted by their fold-change and any adjacent ChIP binding sites were identified. We performed a Kolmogorov-Smirnov (KS) test comparing the expression-sorted ChIP binding presence vs. a uniform distribution of binding sites, similar to Gene Set Enrichment Analysis [61]. If ChIP binding significantly correlates with the gene expression fold-change, the KS statistic, *d*, will also have significant, non-zero magnitude. To better visualize the KS test, we plotted the difference between the presence of cohesin binding at (expression-sorted) genes in Figure 5. The x axis of this figure is the (fold-change-based) gene rank, and the y axis is the KS statistic *d*, which behaves like a running enrichment score and is higher (lower) when binding sites co-occur more (less) often than expected if there were no correlation between ChIP binding and expression fold-change. The KS test uses only the *d* with the highest magnitude, which is indicated in the plots by a vertical red line. To better visualize ChIP binding presence, we further plot an x-mirrored density of peak presence at the top of each plot; the gray "beanplot" [62] at the top of the plots are larger when many of the genes have adjacent ChIP binding sites.

### siRNA depletion

Wild type MEFs were transfected using HiPerFect (Qiagen) following the manufacturer’s protocol with 10mM siRNA. A mixture of 30μl HiPerFect, 3μl of 20μM siRNA, and 150μl DMEM was incubated for 10 mins and added to 2 × 10^6^ cells in 4 ml DMEM. After 6 hrs, 4 ml fresh DMEM with 10% FBS was added. Transfection was repeated the next day. Cells were harvested 48 hrs after the first transfection. SiRNAs against *Nipbl* (Nipbl-1: 5’-GTGGTCGTTACCGAAACCGAA-3’; Nipbl-2: 5’-AAGGCAGTACTTAGACTTTAA-3’) and *Rad21* (5’-CTCGAGAATGGTAATTGTATA-3’) were made by Qiagen. AllStars Negative Control siRNA was obtained from Qiagen.

### RT-q-PCR

Total RNA was extracted using the Qiagen RNeasy Plus kit. First-strand cDNA synthesis was performed with SuperScript II (Invitrogen). Q-PCR was performed using the iCycler iQ Real-time PCR detection system (Bio-Rad) with iQ SYBR Green Supermix (Bio-Rad). Values were generated based on Ct and normalized to control gene Rnh1. PCR primers specific for major satellite, minor satellite, rDNA, and SINE B1 were previously described [48]. Other unique primers are listed in Table 1. The RT-qPCR analyses of the wild type and mutant cells were done with two biological replicates with consistent results. The gene expression changes after siRNA treatment were evaluated with two to three biological replicates with similar results.

### 3C analysis

The chromosome conformation capture (3C) protocol was performed as described [35]. Approximately 1 × 10^7^ cells were crosslinked with 1% formaldehyde at 37°C for 10 mins. Crosslinking was stopped by adding glycine to a final concentration of 0.125M. Cells were centrifuged and lysed on ice for 10 minutes. Nuclei were washed with 500μl of 1.2x restriction enzyme buffer and resuspended with another 500μl of 1.2x restriction enzyme buffer with 0.3% SDS and incubated at 37°C for 1 hr. Triton X-100 was added to 2% and incubated for another 1 hr. 800 U of restriction enzyme (HindIII New England Biolabs) was added and incubated overnight at 37°C. The digestion was heat-inactivated the next day with 1.6% SDS at 65°C for 25 minutes. The digested nuclei were added into a 7ml 1x ligation buffer with 1% Triton X-100, followed by 1 hour incubation at 37°C. T4 DNA ligase (2000 U) (New England Biolabs) was added and incubated for 4 hrs at 16°C followed by 30 minutes at room temperature. Proteinase K (300μg) was added and the sample was reverse-crosslinked at 65°C overnight. Qiagen Gel Purification Kits were used to purify DNA. Approximately 250ng of template was used for each PCR reaction. PCR products were run on 2% agarose gels with SYBRSafe (Invitrogen), visualized on a Fujifilm LAS-4000 imaging system and quantified using Multigauge (Fujifilm).

To calculate interaction frequencies, 3C products were normalized to the constitutive interaction at the excision repair cross-complementing rodent repair deficiency, complementation group 3 (ercc3) locus [63, 64], which is unaffected in mutant MEFs. A control template was made to control for primer efficiencies locus-wide as described [65]. PCR fragments spanning the restriction sites examined were gel purified and equimolar amounts were mixed (roughly 15μg total) and digested with 600 U restriction enzyme overnight and subsequently ligated at a high DNA concentration (>300ng/μl). The template was purified with the Qiagen PCR Purification Kit and mixed with an equal amount of digested and ligated genomic DNA. 250ng of the resulting control template was used for each PCR for normalization against PCR primer efficiencies. Two biological replicates with three technical replicates each were analyzed for both wild type and mutant cells and for control and Nipbl siRNA-treated cells, which yielded consistent results.

## RESULTS

### Nipbl haploinsufficiency leads to a global reduction of cohesin binding to its binding sites

In order to investigate how Nipbl haploinsufficiency leads to CdLS, cohesin binding was examined genome-wide by ChIP-seq analyses using antibody specific for the cohesin subunit Rad21, in wild type and *Nipbl* +/- mutant MEFs derived from E15.5 embryos [19] (Fig. 1A). MEFs derived from five wild type and five mutant pups from two litters were combined to obtain sufficient chromatin samples for ChIP-seq analysis. *Nipbl* +/- mutant MEFs express approximately 30-40% less Nipbl compared to wild type MEFs [19] (Table 2). MEFs from this embryonic stage were chosen in order to match with a previous expression microarray study, because they are relatively free of secondary effects caused by *Nipbl* mutation-induced developmental abnormalities compared to embryonic tissue [19]. Consistent with this, there is no noticeable difference in growth rate and cell morphology between normal and mutant MEFs [19]. This particular anti-Rad21 antibody was used previously for ChIP analysis and was shown to identify holo-cohesin complex binding sites [30, 35, 45, 66]. This is consistent with the close correlation of the presence of other cohesin subunits at identified Rad21 binding sites [67] (Fig. 1B).

**Figure 1.**
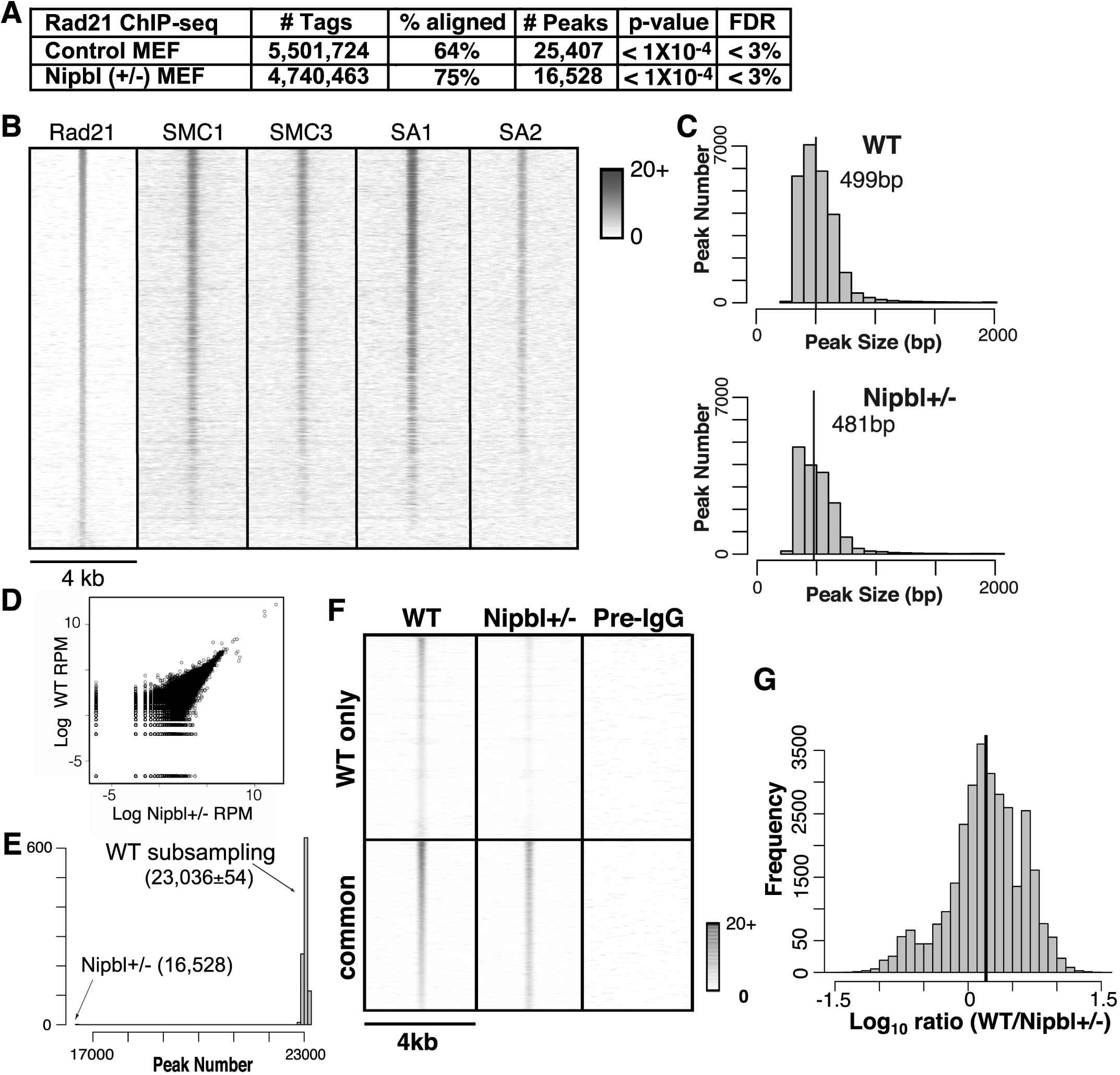
Global decrease of cohesin binding to chromatin in *Nipbl* heterozygous mutant MEFs. **A.** Cohesin binding sites identified by ChIP-sequencing using antibody specific for Rad21 in control wild type and Nipbl +/- MEFs. Peak calling was done using AREM [50]. The *p*-value and FDR are shown. **B.** Heatmap comparison of Rad21 ChIP-seq data with those of SMC1, SMC3, SA1 and SA2. Rad21 peaks in the wild type MEFs are ranked by strongest to weakest, and compared to the ChIP-seq data of SMC1, SMC3, SA1 and SA2 in MEFs (GSE32320) [67] in the corresponding regions. The normalized (reads per million) tag densities in a 4 kb window around each Rad21 peak are plotted, with peaks sorted from the highest number of tags in the wild type MEFs to the lowest. **C.** Histogram of cohesin peak widths in wild type and mutant MEFs, indicating the number of peaks in a given size range. The segmentation of the histogram is at 100bp intervals. The median value is indicated with a vertical black line and labeled. **D.** Scatter plot of histone H3 ChIP-seq tag counts in wild type and mutant MEFs in 500 bp bins across the mouse genome. The values are plotted in log reads per million (RPM). **E.** Histogram showing the distribution of total peaks called. A comparable number of reads to the *Nipbl*+/- mutant dataset (i.e. 4,740,463) were sub-sampled from the wild type dataset, and peaks called using only the sub-sampled reads. This process was performed 1000 times to produce the histogram above. Mean values with standard deviations are shown. **F.** Heatmap analysis of cohesin binding in wild type (WT) MEFs and corresponding peak signals in *Nipbl* +/- MEFs. The normalized (reads per million) tag densities in a 4 kb window around each peak are plotted, with peaks sorted from the highest number of tags in the wild type to the lowest. Peaks are separated into two categories, those that are found only in wild type (“WT only”) and those that overlap between wild type and *Nipbl* +/- (“common”). Preimmune IgG ChIP-seq signals in the corresponding regions are also shown as a control. The color scale indicates the number of tags in a given region. **G.** Histogram of the ratio between normalized (reads per million total reads) wild type and mutant reads in peaks common to both. Positive values indicate more wild type tags. The black line indicates the mean ratio between wild type and mutant tag counts.

**Table 2.** Nipbl and Rad21 depletion levels in mutant and siRNA-treated MEFs.

| Gene | Nipbl+/- mutant | Nipbl siRNA | Rad21 siRNA |
| --- | --- | --- | --- |
| Nipbl | 0.68±0.003 | 0.68±0.001 | 1.04±0.051 |
| Rad21 | 0.94±0.021 | 0.99±0.021 | 0.26±0.018 |
| CTCF | 0.95±0.050 | 0.96±0.066 | 0.84±0.074 |

Cohesin binding sites were identified using AREM [50], with a significance cut-off based on a *p*-value less than 1 × 10^−4^, resulting in a FDR below 3.0% (Fig. 1A). Cohesin binding peaks ranged from ∼200bp to ∼6kb in size with the majority less than 1kb in both wild type and mutant cells (median value of 499 bp in wild type and 481 bp in mutant cells) (Fig. 1C). Approximately 35% fewer cohesin binding sites were found in *Nipbl* +/- mutant MEFs compared to the wild type MEFs (Fig. 1A). This is not due to variability in sample preparation since no significant difference in the histone H3 ChIP-seq was observed between the wild type and mutant cell samples (R-value=0.96) (Fig. 1D). Since the total read number for mutant ChIP-seq was ∼15% less than for wild type ChIP-seq (Fig. 1A), we examined whether the difference was in part due to a difference in the number of total read sequences between the two Rad21 ChIP samples. To address this, we randomly removed reads from the wild type sample to match the number of reads in the mutant sample, and ran the peak discovery algorithm again on the reduced wild type read set. This was repeated 1,000 times. We found that the wild type sample still yielded ∼39% more peaks than the mutant, indicating that identification of more peaks in the wild type sample is not due to a difference in the numbers of total read sequences (Fig. 1E). Thus, cohesin appears to bind to fewer binding sites in *Nipbl* haploinsufficient cells.

The above results might suggest that a significant number of binding sites are unique to the wild type cells (Fig. 1A). When we compared the raw number of reads located within wild type peaks and the corresponding regions in mutant MEFs, however, we noted a reduced, rather than a complete absence of, cohesin binding in mutant cells (Fig. 1F). Those regions in mutant cells corresponding to the “WT only” regions consistently contain one to three tags in a given window, which are below the peak cut-off. However, the signals are significant compared to the negative control of preimmune IgG (Fig. 1F). Furthermore, even for those sites that are apparently common between the control and mutant MEFs, the binding signals appear to be weaker in mutant cells (Fig. 1F). To validate this observation, we segmented the genome into nonoverlapping 100bp bins, and plotted a histogram of the log ratios of read counts between the wild type and mutant samples in each bin, with read counts normalized using reads per kb per million total reads (RPKM) [68]. The plot indicates that the read counts for the mutant bins are generally less than those for the wild type bins, even for the binding sites common to both wild type and mutant cells (Fig. 1G). Signal intensity profiles of the Rad21 ChIP-seq in the selected gene regions also show a general decrease of Rad21 binding at its binding sites in Nipbl+/- MEFs compared to the control MEFs (see below, Fig. 6B). Decreased cohesin binding was further confirmed by manual ChIP-qPCR analysis of individual cohesin binding sites using at least three independent control and mutant MEF samples supporting the reproducibility of the results (see below, Figure 3). Decreased cohesin binding was also observed at additional specific genomic regions in *Nipbl+/-* MEFs [69]. Taken together, the results indicate that cohesin binding is generally decreased at its binding sites found in wild type MEFs, rather than re-distributed, in mutant MEFs.

### The relationship of cohesin binding sites with CTCF binding sites and CTCF motifs

It has been reported that cohesin binding significantly overlaps with CTCF sites and depends on CTCF [30, 31]. A study in mouse embryonic stem cells (mESCs) showed, however, that there is only a limited overlap between CTCF- and Nipbl-bound cohesin sites, suggesting that there are two categories of cohesin binding sites, and the latter may be particularly important for gene activation [40]. Other studies also revealed that ∼20-30% of cohesin sites in different human cancer cell lines and up to ∼50% of cohesin sites in mouse liver appear to be CTCF-free [42, 43]. Some of these non-CTCF sites overlap with sequence-specific transcription factor binding sites in a cell type-specific manner, highlighting the apparent significance of CTCF-free cohesin sites in cell type-specific gene expression [42, 43]. *De novo* motif discovery by MEME identified the CTCF motif to be the only significant motif associated with cohesin binding sites in our MEFs (Fig. 2A). Comparing our cohesin peaks with experimentally determined CTCF binding peaks in MEFs [40], we found that approximately two-thirds of cohesin binding sites detected by Rad21 ChIP overlapped CTCF binding sites (Fig. 2B). This is comparable with what was initially observed in mouse lymphocytes [30] and HeLa cells [31] using antibodies against multiple cohesin subunits. In contrast to recent studies reporting that almost all the CTCF binding sites overlap with cohesin [43], our results show that less than 60% of CTCF binding sites are co-occupied with cohesin (Fig. 2B). This is consistent with the fact that CTCF binds and functions independently of cohesin at certain genomic regions [34, 41, 70, 71].

**Figure 2.**
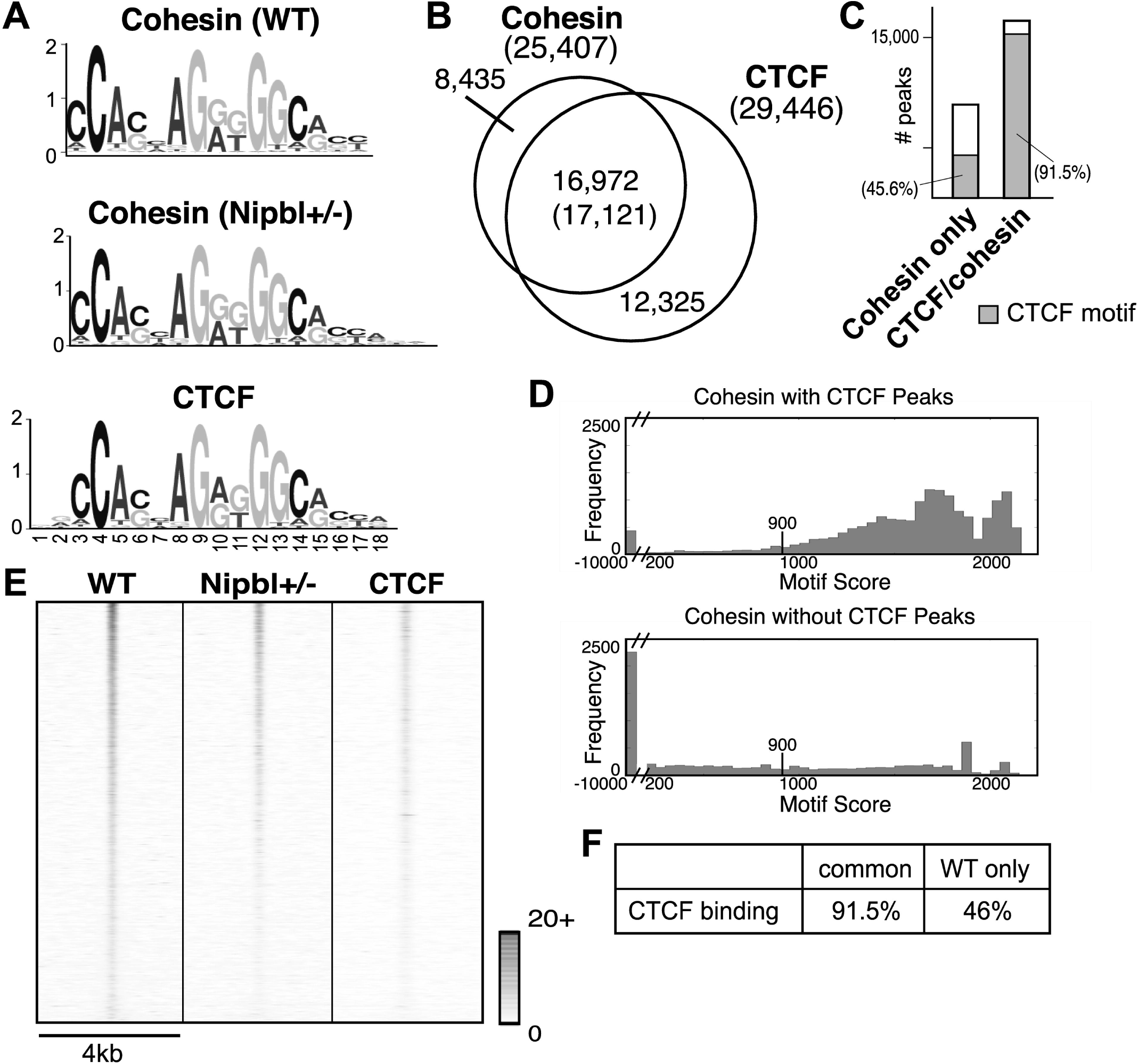
Most of cohesin binding sites contain CTCF motifs. **A.** De novo motif search of cohesin binding sites using MEME. The CTCF motifs identified at the cohesin binding sites in WT and mutant MEFs are compared to the CTCF motif obtained from CTCF ChIP-seq data in MEFs (GSE22562) [40]. E-values are 5.5e-1528 (cohesin binding sites in WT MEFs), 6.6e-1493 (cohesin binding sites in Nipbl MEFs), and 2.6e-1946 (CTCF binding sites in MEFs), respectively. **B.** Overlap of cohesin binding sites with CTCF binding sites. The number in the parenthesis in overlapping regions between cohesin and CTCF binding represents the number of CTCF binding peaks. **C.** Presence of CTCF motifs in cohesin only and cohesin/CTCF binding sites. Shaded area represents binding sites containing CTCF motifs defined in (A) (FDR 4.7%). **D.** The CTCF motif score distribution for all cohesin peaks that overlap with a CTCF peak (top) and that don't overlap with a CTCF peak (bottom). Note that the X axis is discontinuous and scores less than 200 are placed in the single bin in each figure. For peaks that contained multiple CTCF motifs, we report the maximum score for the peak. The score threshold (900 with FDR 4.7%) is marked in each figure. **E.** Heatmap comparison of cohesin ChIP-seq tags in WT MEFs and Nipbl mutant MEFs with CTCF ChIP-seq tags at the corresponding regions in wild type MEFs [40] as indicated at the top. The normalized (reads per million total reads) tag densities in a 4 kb window (±2kb around the center of all the cohesin peaks) are plotted, with peaks sorted by the number of cohesin tags (highest at the top) in WT MEFs. Tag density scale from 0 to 20 is shown. **F.** Percentages of CTCF binding in cohesin binding sites common or unique to WT MEFs.

Presence of a CTCF motif closely correlates with CTCF binding: over 90% of cohesin binding sites overlapping with CTCF peaks contain CTCF motifs (Fig. 2C). In contrast, less than half of cohesin binding sites harbor CTCF motifs in the absence of CTCF binding. Cohesin binding sites without CTCF binding tend to be highly deviated from a CTCF motif, reflecting a CTCF-independent mechanism of recruitment (Fig. 2D). Interestingly, a small population of cohesin-CTCF overlapping sites that also lack any CTCF motif, suggesting an alternative way by which cohesin and CTCF bind to these regions (Fig. 2C and D).

### Nipbl reduction affects cohesin binding at CTCF-bound sites and repeat regions

In mESCs, it was proposed that Nipbl and CTCF recruit cohesin to different genomic regions, implying that cohesin binding to CTCF sites may be Nipbl-independent [40]. We noticed that when we ranked cohesin binding sites based on the read number in wild type peaks, they matched closely with the ranking of cohesin binding sites in mutant MEFs, indicating that the decrease of cohesin binding is roughly proportional to the strength of the original binding signals (Fig. 2E). This suggests that most cohesin binding sites have similar sensitivity to Nipbl reduction. Importantly, CTCF binding signals also correlate with the ranking of cohesin binding, indicating that CTCF-bound sites are in general better binding sites for cohesin (Fig. 2E). Because of this, they satisfy the peak definition despite the decrease of cohesin binding in mutant cells (Fig. 1F and G, and Fig. 6B). This explains why CTCF-bound cohesin sites are apparently enriched in the sites that are common to both wild type and mutant cells (Fig. 2F).

Based on the above data, we further clarified the role of Nipbl in cohesin binding to CTCF sites. We compared the effect of Nipbl reduction on cohesin binding to representative sites, which have either CTCF binding or a CTCF motif or both (Fig. 3A). Decreased cohesin binding was observed at sites tested by manual ChIP-qPCR in Nipbl mutant MEFs, correlating with the decreased Nipbl binding (Fig. 3A). Consistent with the genome-wide ChIP-seq analysis (Fig. 1D), control histone H3 ChIP-qPCR revealed no significant differences at the corresponding regions, indicating that the decreased cohesin binding is not due to generally decreased ChIP efficiency in mutant MEFs compared to the wild type MEFs (Fig. 3A, bottom). Similar results were obtained using a small interfering RNA (siRNA) specific for *Nipbl* (Fig. 3C), which reduced Nipbl to a comparable level as in mutant cells (western blot in Fig. 3B and RT-qPCR results in Table 2). This demonstrates the specificity of the Nipbl antibody and confirms that the decreased cohesin binding seen in Nipbl mutant MEFs is the consequence of reduced Nipbl (Fig. 3A). Thus, Nipbl also functions in cohesin loading at CTCF sites.

**Figure 3.**
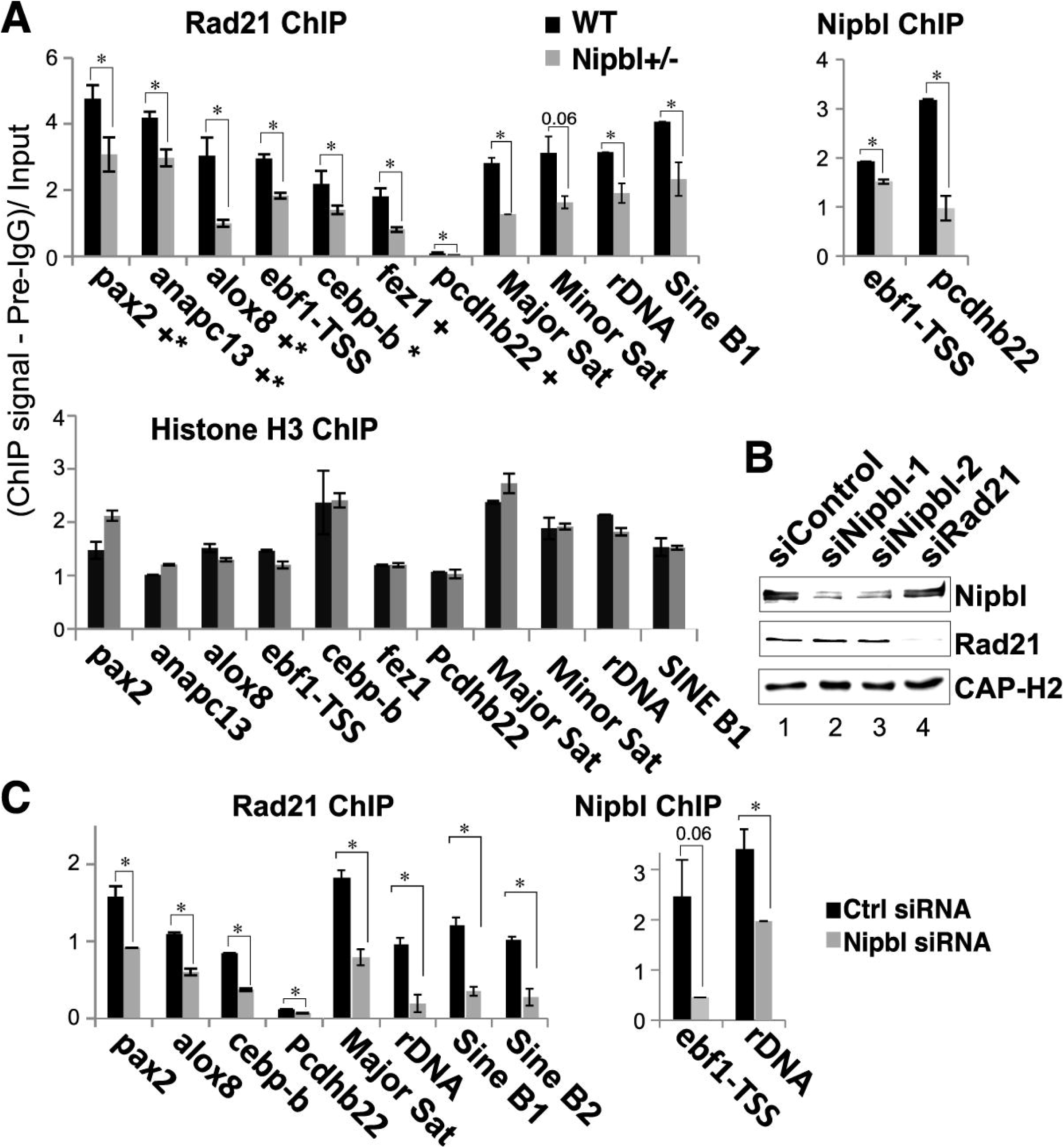
Nipbl reduction decreases cohesin binding. **A.** Manual ChIP-q-PCR of cohesin binding sites at unique gene regions and repeat regions using anti-Rad21 antibody (top left) compared to histone H3 (bottom) in Nipbl +/- mutant and wild type MEFs. Representative examples of Nipbl ChIP are also shown (top, right). “+” indicates CTCF binding and “*” indicates the presence of motif. PCR signals were normalized with preimmune IgG (pre-IgG) and input. *p<0.05. **B.** Western blot analysis of control, Nipbl or Rad21 siRNA-treated cells is shown using antibodies indicated. Depletion efficiency and specificity of Nipbl siRNA were also examined by RT-q-PCR (Table 2). Comparable depletion efficiencies and ChIP results were obtained by two Nipbl siRNA (siNipbl-1 and siNipbl-2) (lanes 2 and 3; data not shown). **C.** Similar manual ChIP-q-PCR analysis as in (A) in control and Nipbl siRNA (siNipbl-1)-treated MEFs.

Repeat sequences are often excluded from ChIP-seq analysis. However, cohesin binding is found at various repeat sequences, including pericentromeric and subtelomeric heterochromatin, and ribosomal DNA regions in the context of heterochromatin in mammalian cells [44, 45]. Thus, we also tested the effect of Nipbl reduction on cohesin binding to repeat sequences by manual ChIP-PCR (Fig. 3A). Both *Nipbl* mutation (Fig. 3A, top) and Nipbl depletion by siRNA (Fig. 3B and C) resulted in decreased cohesin binding at the repeat regions, indicating that Nipbl is also important for cohesin binding to repeat sequences. In contrast, there were no significant differences in the histone H3 ChIP signals between these repeat regions in wild type and mutant MEFs (Fig. 3A, bottom). Taken together, the results indicate that Nipbl functions in cohesin loading even at CTCF sites and repeat regions, confirming the genome-wide decrease of cohesin binding caused by Nipbl haploinsufficiency.

### Cohesin distribution patterns in the genome and enrichment in promoter regions

In order to gain insight into how the weakening of cohesin binding may affect gene expression in mutant cells, the distribution of cohesin binding sites in the genomes of both wild type and mutant MEFs were examined. Approximately 50% of all cohesin binding sites are located in intergenic regions away from any known genes (Fig. 4A). However, there is a significant enrichment of cohesin binding in promoter regions, and to a lesser extent in the 3’ downstream regions, relative to the random genomic distribution generated by sampling from pre-immune ChIP-seq reads (Fig. 4B). Similar promoter and downstream enrichment has been observed in mouse and human cells [30, 31, 40, 42, 67] as well as in Drosophila [72]. Promoter enrichment is comparable in both wild type and *Nipbl* mutant MEFs, constituting ∼10% of all the cohesin binding sites (Fig. 4A). Thus, there is no significant redistribution or genomic region-biased loss of cohesin binding sites in Nipbl mutant cells.

**Figure 4.**
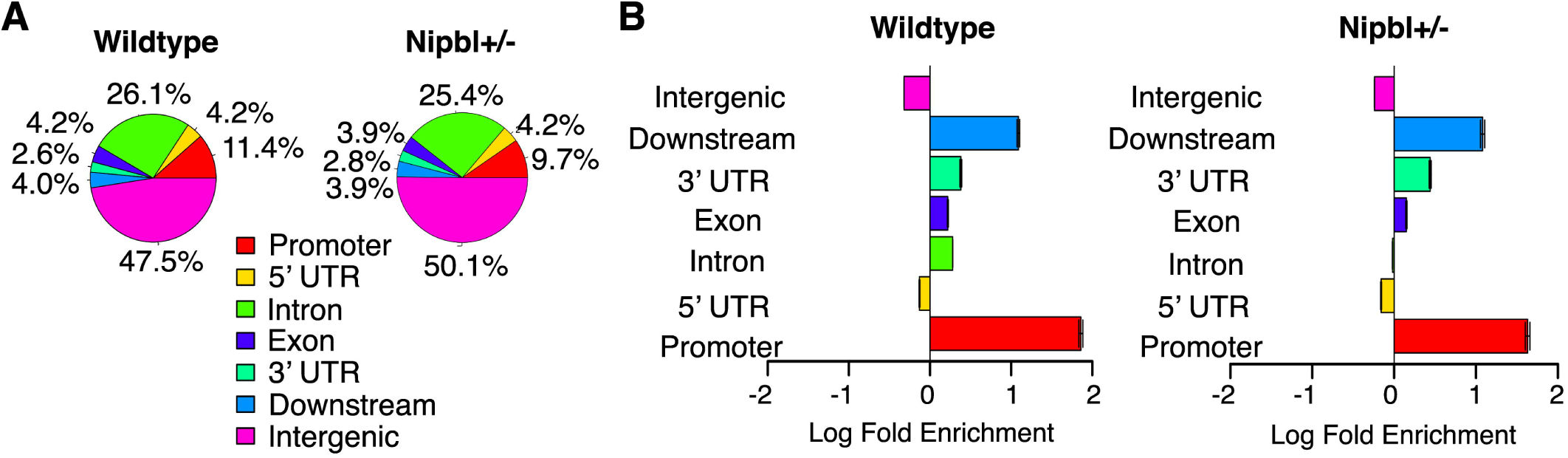
Cohesin binding site distribution in the genome in MEFs. **A.** Percentage distribution of cohesin peaks in genomic regions. “Promoter” and “Downstream” is defined as 2500bp upstream of the transcription start site (TSS) and 500 bp downstream of the TSS, and “Downstream” represents 500 bp upstream of transcription termination site (TTS) and 2500 bp downstream of TTS. The 3’ and 5’ untranslated regions (UTRs) are defined as those annotated by the UCSC genome browser minus the 500 bp interior at either the TSS or TTS. When a peak overlaps with multiple regions, it is assigned to one region with the order of precedence of promoter, 5’ UTR, Intron, Exon, 3’UTR, downstream, and intergenic. **B.** Enrichment of cohesin peaks across genomic regions as compared to randomly sampled genomic sequence. A comparable number of peaks (25,407 and 16,528 peaks in wild type and mutant MEFs, respectively), with the same length as the input set, were randomly chosen 1000 times and the average used as a baseline to determine enrichment in each genomic region category.

### Cohesin-bound genes are sensitive to Nipbl haploinsufficiency

Based on the significant enrichment of cohesin binding in the promoter regions, we next examined the correlation between cohesin binding to the gene regions and the change of gene expression in mutant MEFs using a KS test. This is a nonparametric test for comparing peak binding sites with gene expression changes in the mutant MEFs (Fig. 5). Genes that displayed the greatest expression change in mutant MEFs compared to the wild type MEFs showed a strong correlation with cohesin binding to the gene region, indicating that direct binding to the target genes is the major mechanism by which cohesin mediates gene regulation in a Nipbl dosage-sensitive fashion (Fig. 5A, left). Random sampling of a comparable number of simulated peaks in the gene regions yielded no correlation (Fig. 5D, left). Interestingly, cohesin binding to the gene region correlates better with decreased gene expression than increased expression in mutant cells, indicating that gene activation, rather than repression, is the major mode of cohesin function at the gene regions (Fig. 5A, middle).

**Figure 5.**
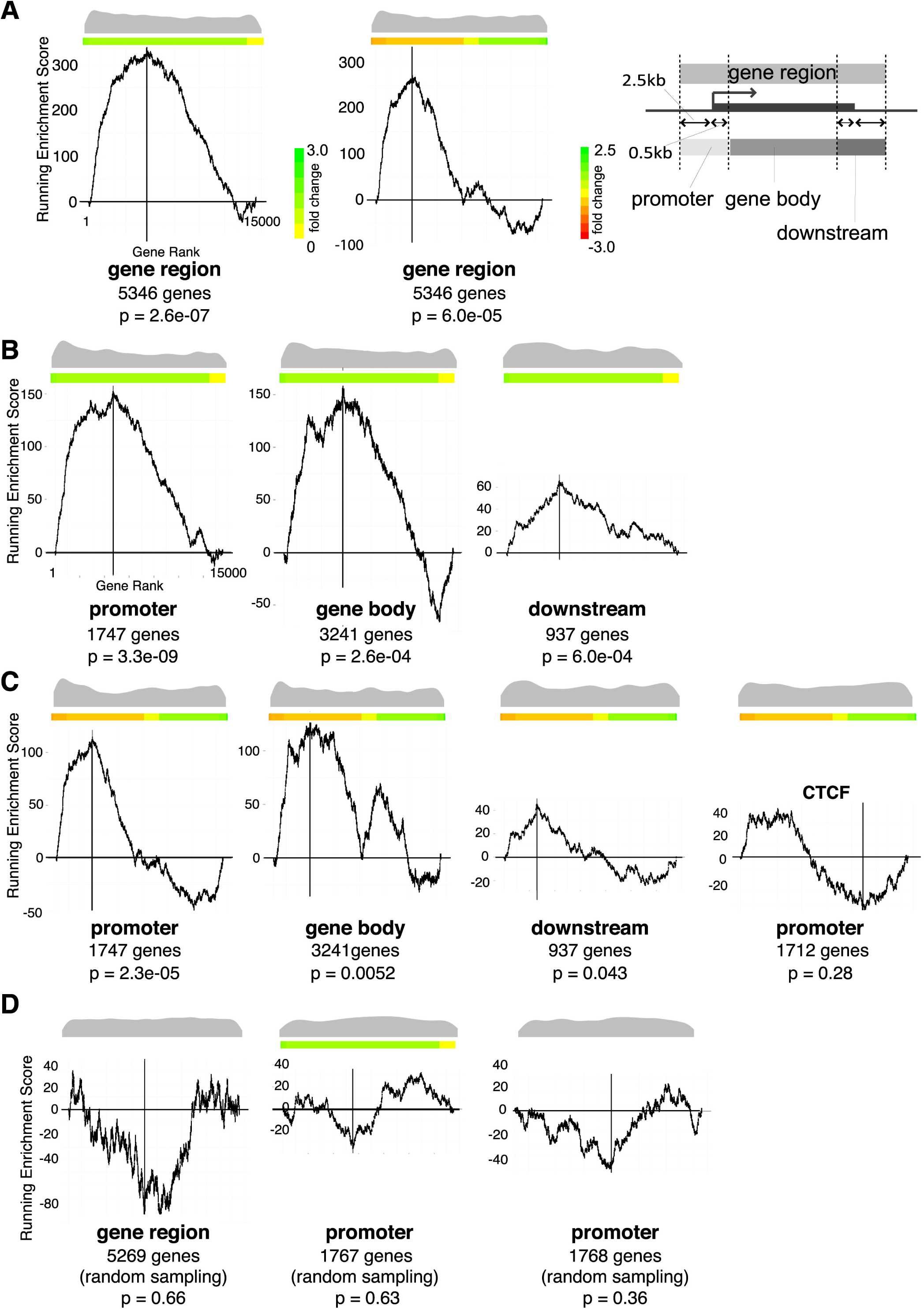
Correlation of cohesin binding and gene expression changes in mutant MEFs. **A.** KS test indicating the degree of cohesin binding to genes changing expression in *Nipbl* +/- MEFs. X-axis represents all 13,587 genes from the microarray data [19] ranked by absolute fold expression changes from biggest on the left to the smallest on the right in the left panel. Fold changes are shown in different colors as indicated on the side. In the middle panel, gene expression changes were ranked from negative to positive with the color scale shown on the side. Both color scales apply to the rest of the Figure. The Y-axis is the running enrichment score for cohesin binding (see EXPERIMENTAL PROCEDURES for details). Distribution of cohesin-bound genes among 13,587 genes examined is shown as a beanplot [62] at the top, and the number of cohesin-bound genes and *p*-values are shown underneath. The schematic diagram showing the definition of the gene regions, promoter (2.5kb upstream and 0.5kb downstream of TSS), gene body, and downstream (2.5kb downstream and 0.5kb upstream of TTS) regions is shown on the right. **B.** Similar KS test analysis as in (A), in which cohesin binding to the promoter, gene body, and downstream regions are analyzed separately. **C.** Genes are ranked by expression changes from positive on the left to negative on the right. Fold changes are shown by different colors as indicated on the right. CTCF binding to promoter regions (GSE22562) [40] was analyzed for a comparison. **D.** Lack of correlation between the mutant expression changes and randomly chosen genes are shown on the right as a negative control.

When analyzed separately, cohesin binding to the promoter regions (+2.5kb to -0.5kb of transcription start sites (TSS) (Fig. 5A, right)) showed the highest correlation (*p*-value=3.3e-09) compared to the gene body and downstream (Fig. 5B). Thus, cohesin binding to the promoter regions is most critical for gene regulation. Similar to the entire gene region, cohesin binding correlates more significantly with a decrease in gene expression in mutant cells, which is particularly prominent at the promoter regions compared to gene bodies or downstream, indicating the significance of cohesin binding to the promoter regions in gene activation (Fig. 5C). Although cohesin and CTCF binding closely overlapped at promoter regions in HeLa cells [31], the overlap of CTCF binding with cohesin in MEFs is lower in the promoter regions (54%) than that in the intergenic regions (67%) [40]. Consistent with this, there is no significant correlation between CTCF binding in the promoter regions and gene expression changes in *Nipbl* mutant MEFs (*p*-value=0.28) by KS test (Fig. 5C, right), further indicating the cohesin-independent and Nipbl-insensitive function of CTCF in gene regulation. Taken together, the results suggest that cohesin binding to gene regions (in particular, to promoters) is significantly associated with gene activation that is sensitive to Nipbl haploinsufficiency.

### Identification of cohesin target genes sensitive to Nipbl haploinsufficiency

The results above indicate that cohesin-bound genes sensitive to a partial loss of Nipbl can be considered to be Nipbl/cohesin target genes. Among 218 genes that changed expression significantly in mutant cells compared to the wild type (>1.2-fold change, *p*-value < 0.05) [19], we found that more than half (115 genes) were bound by cohesin, and thus can be considered Nipbl/cohesin target genes (Table 3). This is a conservative estimate of the number of direct target genes since cohesin binding sites beyond the upstream and downstream cut-offs (2.5 kb) were not considered for the analysis. Consistent with the KS test analysis (Fig. 5), ∼74% of these cohesin target genes were downregulated in mutant cells, indicating that the positive effect of cohesin on gene expression is particularly sensitive to partial reduction of Nipbl (Table 3).

**Table 3.** Gene expression changes and cohesin binding status.

|  | Total | Cohesin binding |  |  |  |  |
| --- | --- | --- | --- | --- | --- | --- |
|  |  | Gene region | Promoter | Gene body | Downstream | None |
| Total | 218 | 115 | 61 | 83 | 20 | 103 |
| Up-regulated | 62 | 30 | 14 | 22 | 6 | 32 |
| Down-regulated | 156 | 85 | 47 | 61 | 14 | 71 |

(Fold change>1.2, *p*-value<0.05)

Many of these Nipbl/cohesin-target genes contain cohesin binding sites in more than one region (promoter, gene body and/or downstream), suggesting their collaborative effects (Fig. 6A). In particular, the promoter binding of cohesin is often accompanied by its binding to the gene body. However, binding pattern analysis revealed no significant correlation between a particular pattern and/or number of cohesin binding sites and gene activation or repression (Fig. 6A). Rad21 ChIP-seq signal intensity profiles of several cohesin target genes (as defined above) reveal decreased cohesin binding in mutant cells at the binding sites originally observed in the wild type cells, supporting the notion that gene expression changes are the direct consequence of the reduced cohesin binding (Fig. 3A; Fig. 6B, top). There are other genes, however, that did not change expression significantly in mutant MEFs, but nevertheless also have reduced cohesin peaks nearby (Fig. 6B, bottom), suggesting that cohesin binding is not the sole determinant of the gene’s expression status and that its effect is context-dependent.

**Figure 6.**
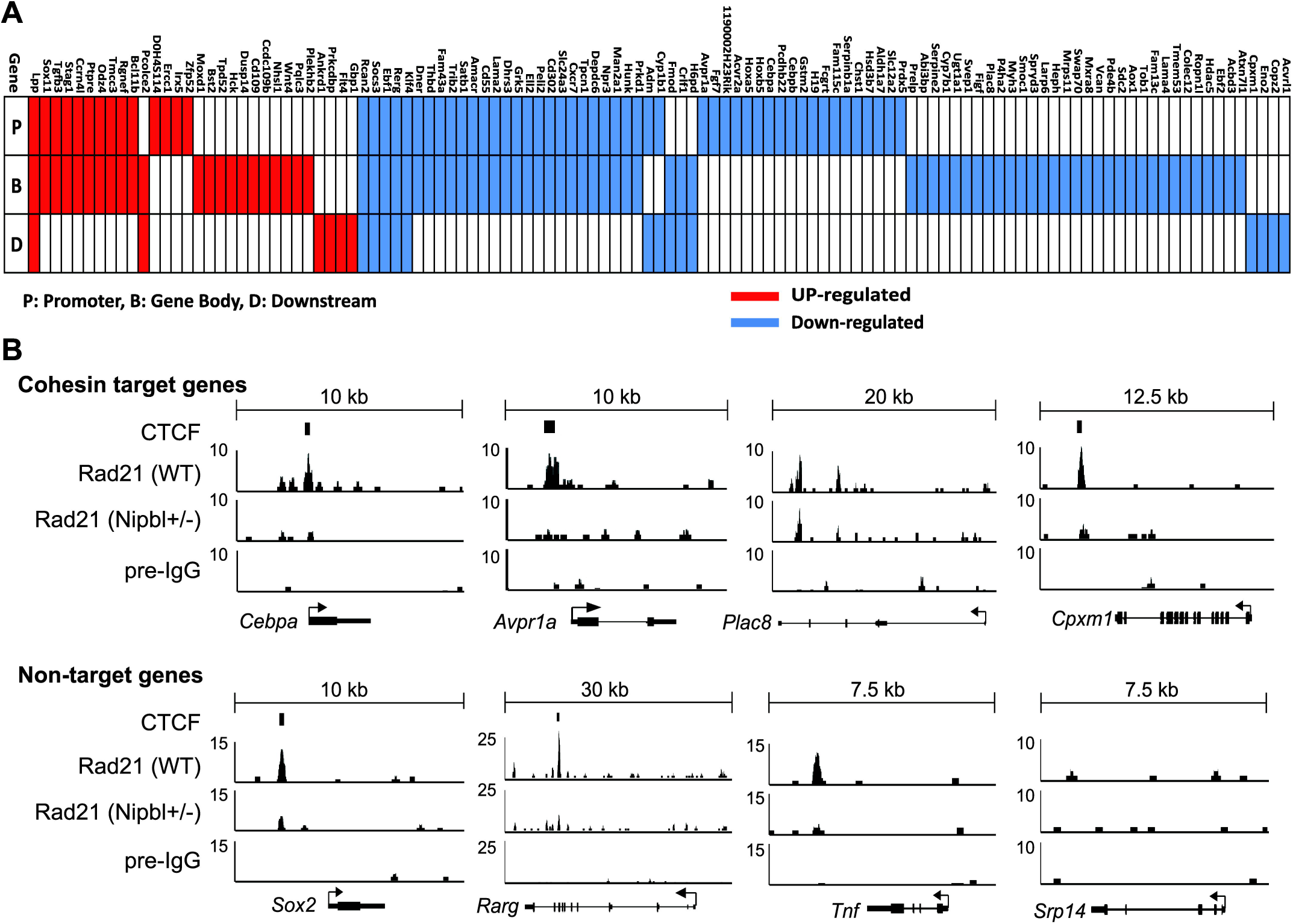
Cohesin binding signals at specific gene regions. **A.** Cohesin binding site distribution in cohesin target genes as defined in Table 1. Cohesin binding to the promoter (P), gene body (B), and/or downstream region (D) are indicated for each cohesin target gene in red (upregulated) and blue (downregulated) boxes. **B.** Signal intensity profiles of Rad21 ChIP-seq at specific gene regions in wild type and Nipbl mutant MEFs. Preimmune IgG ChIP-seq signals are shown as a negative control. Experimentally determined CTCF binding peaks in MEFs [40] are also indicated. Examples of genes that are bound by cohesin and changed expression in Nipbl+/- MEFs (top) and those genes that did not change expression (bottom) are shown. No cohesin binding peaks were found at the *Srp14* gene region.

Gene ontology analysis revealed that the target genes bound by cohesin at the promoter regions and affected by Nipbl deficiency are most significantly enriched for those involved in development (Table 4). The results suggest a direct link between diminished Nipbl/cohesin and the dysregulation of developmental genes, which contributes to the CdLS phenotype.

**Table 4.**
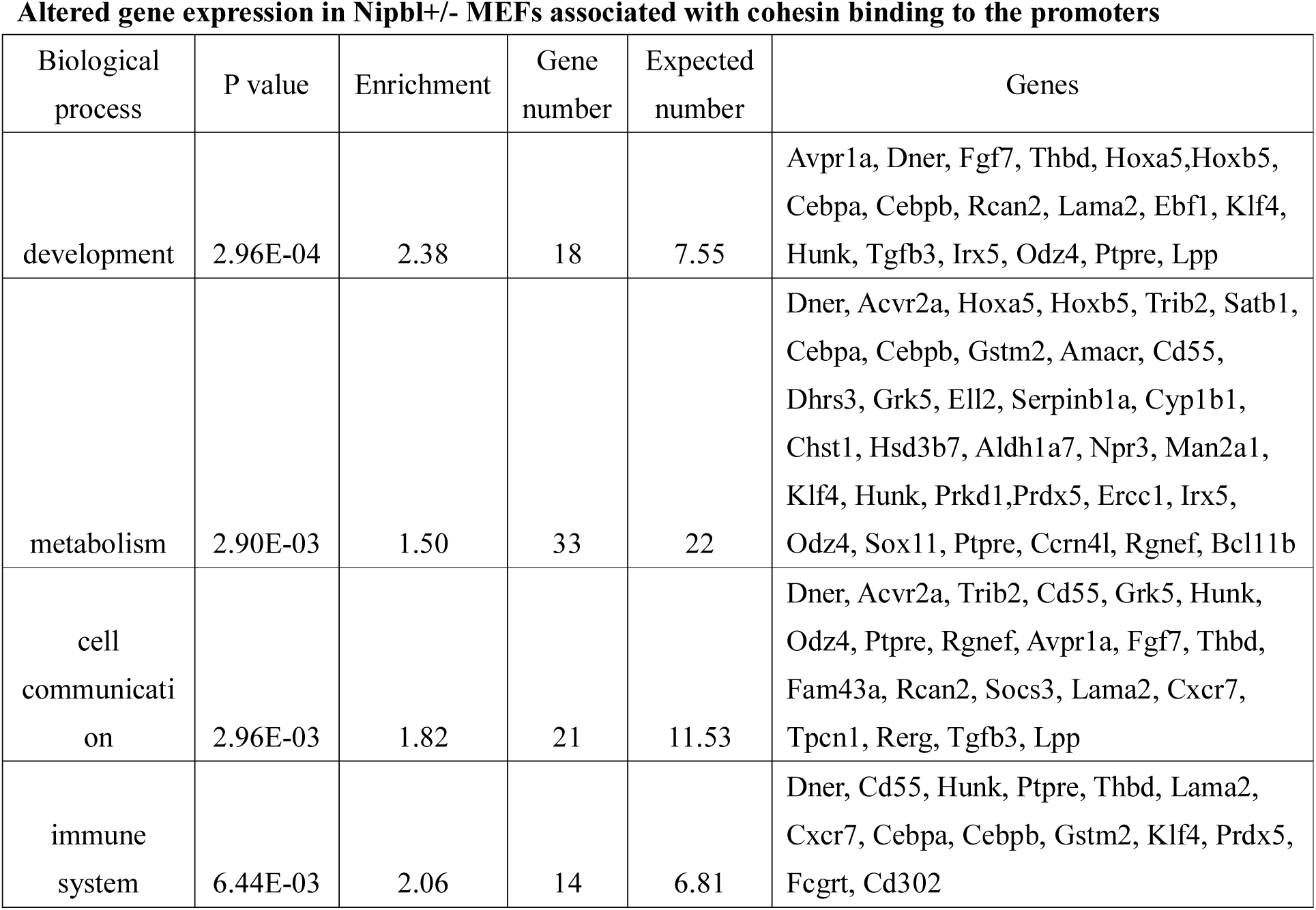
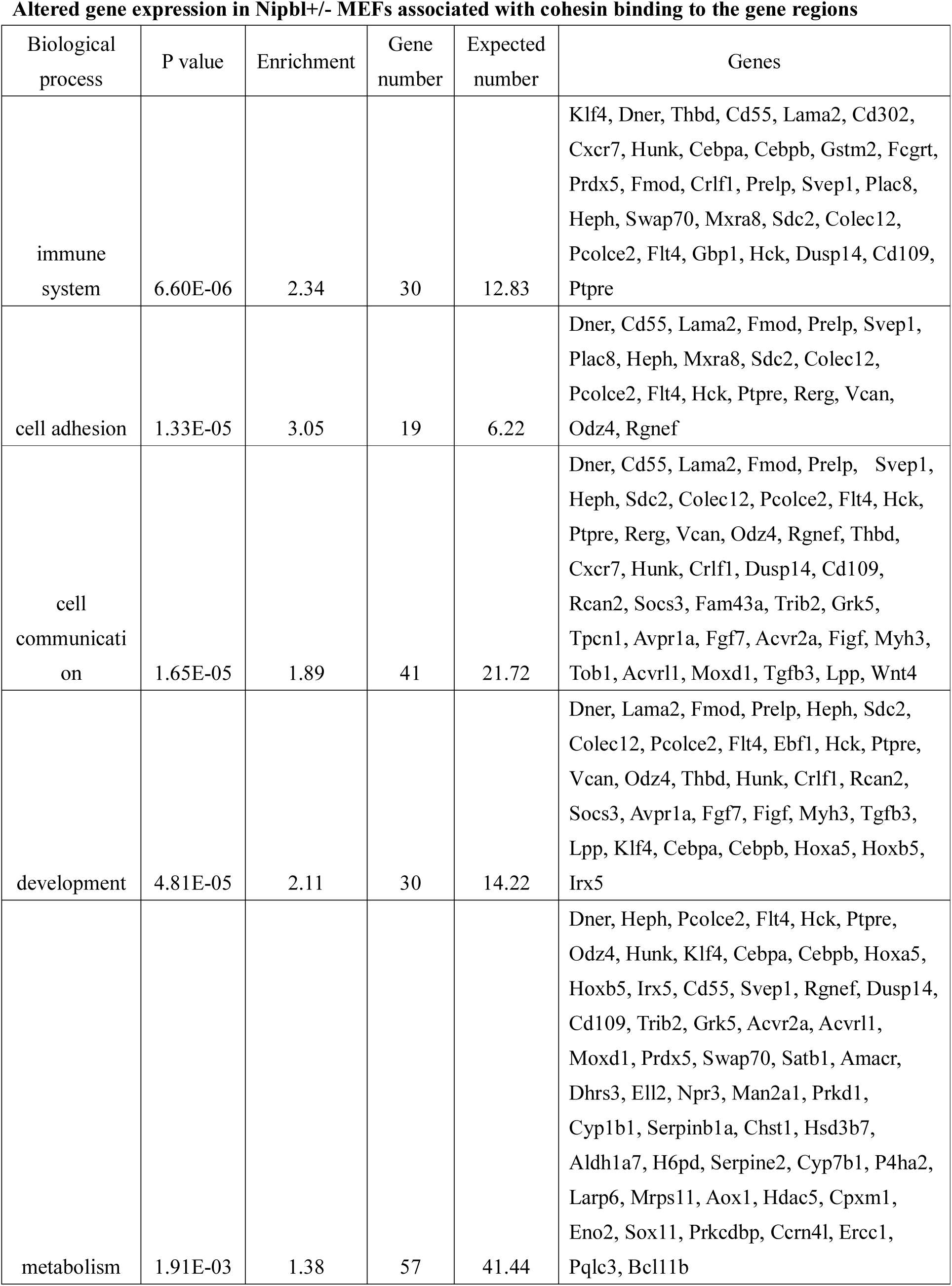
Ontology analysis of cohesin target genes. Biological processes enriched in cohesin target genes with cohesin binding at either promoters or gene regions. “Gene number” is the number of cohesin target genes that belong to a specific category; “Expected number” is the expected gene numbers that belong to a specific category at random.

### Nipbl- and cohesin-mediated activation of adipogenesis genes

One of the reported phenotypes of *Nipbl* +/- mice is their substantial reduction of body fat that mirrors what is observed in CdLS patients [19, 73]. It was found that *Nipbl* +/- MEFs exhibit dysregulated expression of several genes involved in adipocyte differentiation, and reduced spontaneous adipocyte differentiation *in vitro* [19, 73]. We therefore examined the effect of Nipbl haploinsufficiency on these adipogenesis genes in detail. We found that many of them are bound by cohesin, in some cases at multiple sites, suggesting that cohesin plays a direct role in activation of these genes (Fig. 7). Although *Il6* and *Cebpδ* were originally not included in the 115 genes due to low p-values in the microarray analysis (Table 3 and Fig. 6A), significant expression changes were observed in mutant MEFs compared to the wild type MEFs by manual RT-qPCR. *TNFα* and *PPARγ*, involved in adipogenesis, do not change their expression in mutant MEFs [19]. Importantly, a decrease of gene expression was not only observed in *Nipbl* +/- mutant cells, but also by siRNA depletion of Nipbl, confirming that the effect is specifically caused by Nipbl reduction (Fig. 7A). Furthermore, depletion of cohesin itself decreased their expression even more significantly than Nipbl depletion. The results suggest that multiple genes involved in the adipogenesis pathway are direct cohesin targets that are sensitive to Nipbl haploinsufficiency.

**Figure 7.**
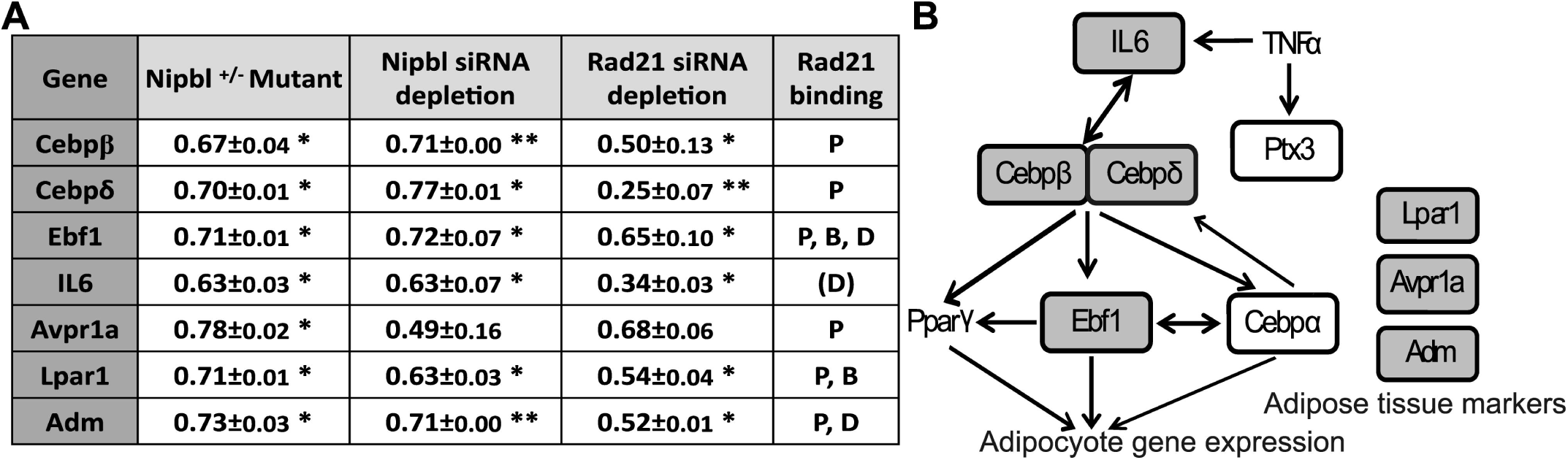
Cohesin plays a direct role in adipogenesis gene regulation. **A.** RT-q-PCR analysis of gene expression changes in *Nipbl* +/- mutant MEFs and MEFs treated with siRNA against *Nipbl* and *Rad21* (* P<0.05, ** P<0.01). Cohesin binding status is also shown. P: promoter, B: gene body, and D: downstream as in Figure 5 with the exception of IL6. For IL6, the cohesin binding site in the downstream region is 3kb away from TSS. **B.** A schematic diagram of genes involved in the adipogenesis pathway. Genes that changed expression in Nipbl +/- mutant MEFs are circled, and those bound by cohesin and examined in (A) are shown with shaded circles.

### Cohesin binding correlates significantly with H3K4me3 at the promoter

To investigate the genomic features associated with cohesin target genes, we examined the chromatin status of the target gene promoters. We found that cohesin peaks closely overlap with the peaks of H3K4me3, a hallmark of an active promoter, in a promoter-specific manner (Fig. 8A). In contrast, there are only minor peaks of H3K27me3 and even less H3K9me3 signal at cohesin-bound promoters, consistent with the results of the KS-test revealing the significant association of cohesin binding to the promoter regions with gene activation rather than repression (Fig. 5C). Interestingly, however, promoter binding of cohesin was found in genes with different expression levels in wild type MEFs, revealing no particular correlation with high gene expression (Fig. 8B). Cohesin target genes defined above (Table 3) also exhibit variable expression levels in wild type MEFs (Fig. 8B). Thus, their expression is altered in Nipbl mutant cells regardless of the original expression level in wild type cells, indicating that cohesin binding contributes to gene expression but does not determine the level of transcription *per se*.

**Figure 8.**
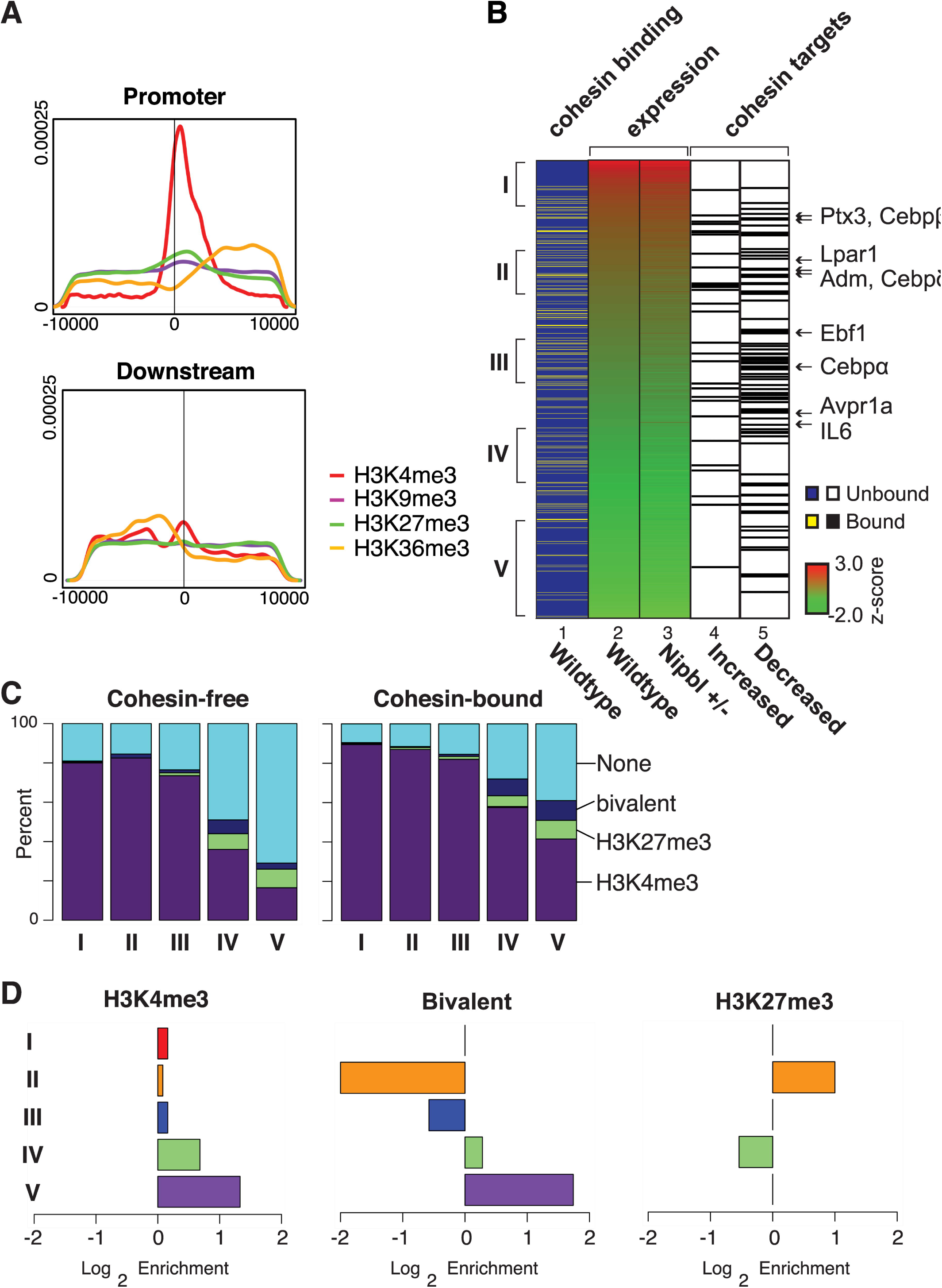
Enrichment of H3K4me3 at the promoters of cohesin-bound genes. **A.** Density of histone modifications within 10kb of cohesin peaks found in the promoter or downstream regions. Histone methylation data was downloaded from NCBI (GEO: GSE26657). Tags within a 10 kb window around cohesin peaks located in a promoter region were counted and normalized to the total number of tags (reads per million) and used to generate a density plot. **B.** Expression status of cohesin target genes. Genes are ranked by their expression status (shown as a z-score) in wild type MEFs (lane 2), and those genes with cohesin binding at the promoter regions are indicated by yellow lines (lane 1). The expression status of the corresponding genes in Nipbl mutant cells is also shown (lane 3), and the cohesin target genes (Table 2) (either upregulated (lane 4) or downregulated (lane 5) in mutant cells) are indicated by black lines. Genes in the adipogenesis pathway are indicated with arrows on the right. Five clusters (I through V) of two hundred cohesin-bound genes each in wild type MEFs according to the expression levels are indicated on the left, which were used for the analysis in (C) and (D). **C.** The numbers of cohesin target genes containing histone marks in the promoter were tallied for the categories I through V from (B). As a control, the cohesin-free gene directly below each cohesin target gene was also tallied and plotted. H3K4me3, H3K9me3, H3K27me3, bivalent (H3K4me3 and H3K27me3), and the promoters with none of these marks (“None”) are indicated. There is almost no signal of H3K9me3 in these categories. **D.** Enrichment plot of H3K4me3, H3K27me3, and bivalent (H3K4me3 and K27me3) in promoters of cohesin-bound genes versus cohesin-free genes in the five expression categories as in (C) is shown.

When cohesin-bound genes were categorized in five different groups based on the gene expression status in wild type MEFs, significant H3K4me3 enrichment was observed even in the cohesin-bound promoters of genes with low expression, compared to cohesin-free promoters of genes with a similar expression level (Fig. 8C). Bivalent (H3K4me3 and H3K27me3) modifications are also enriched in the lowest gene expression category (Fig. 8C). Taken together, the results reveal that there is a close correlation between cohesin binding and H3K4me3 in the promoter regions regardless of the expression levels of the corresponding genes.

### Reduced cohesin binding due to NIPBL reduction can lead to a loss of long-distance chromatin interaction

The above results revealed the critical association of cohesin binding to the promoter region and expression of the target genes. How does cohesin bound to the promoter affect gene expression? We recently showed that cohesin-mediated long-distance chromatin interaction between distal enhancer and promoter regions was reduced at the *β-globin* locus, resulting in reduced gene expression, in *Nipbl* mutant mice [35]. Thus, we tested the potential involvement of cohesin binding to the *Cebpβ* gene, one of the target adipogenesis genes described above, in such long-distance chromatin interaction(s) and whether it is affected by Nipbl reduction using chromosome conformation capture (3C) analysis (Fig. 9). We tested several flanking sites that are positive for cohesin and RNA polymerase II (pol II) binding as well as H3K4me1 and H3K4me3, the hallmarks for enhancers [74-76] (Fig. 9A). We observed that the *Cebpβ* promoter interacts with one such region (Fig. 9A and B, the site “c”). Although the site “c” is associated with only a weak Rad21 ChIP-seq signal, SMC1 and SMC3 ChIP-seq signals were found at the same region [67], confirming that this is an authentic cohesin binding site (Fig. 9A). The results indicate a selectivity of chromatin interactions among neighboring cohesin binding sites, revealing that not all proximal cohesin binding sites interact with each other. Since the other two regions are also bound by CTCF, this may be due to the directionality of CTCF/cohesin binding [77, 78]. Importantly, the observed interaction is indeed reduced in both *Nipbl* mutant and Nipbl siRNA-treated MEFs (Fig. 9B). The 3C signals at the *Cebpβ* locus were normalized to the constant interaction observed at the *Ercc3* locus [63, 64], which was not affected by Nipbl reduction. The results indicate that the decrease of long-distance chromatin interaction involving the promoters and distant DNA elements is one of the direct consequences of reduced cohesin binding, which may be one mechanism of gene expression alteration by Nipbl haploinsufficiency.

**Figure 9.**
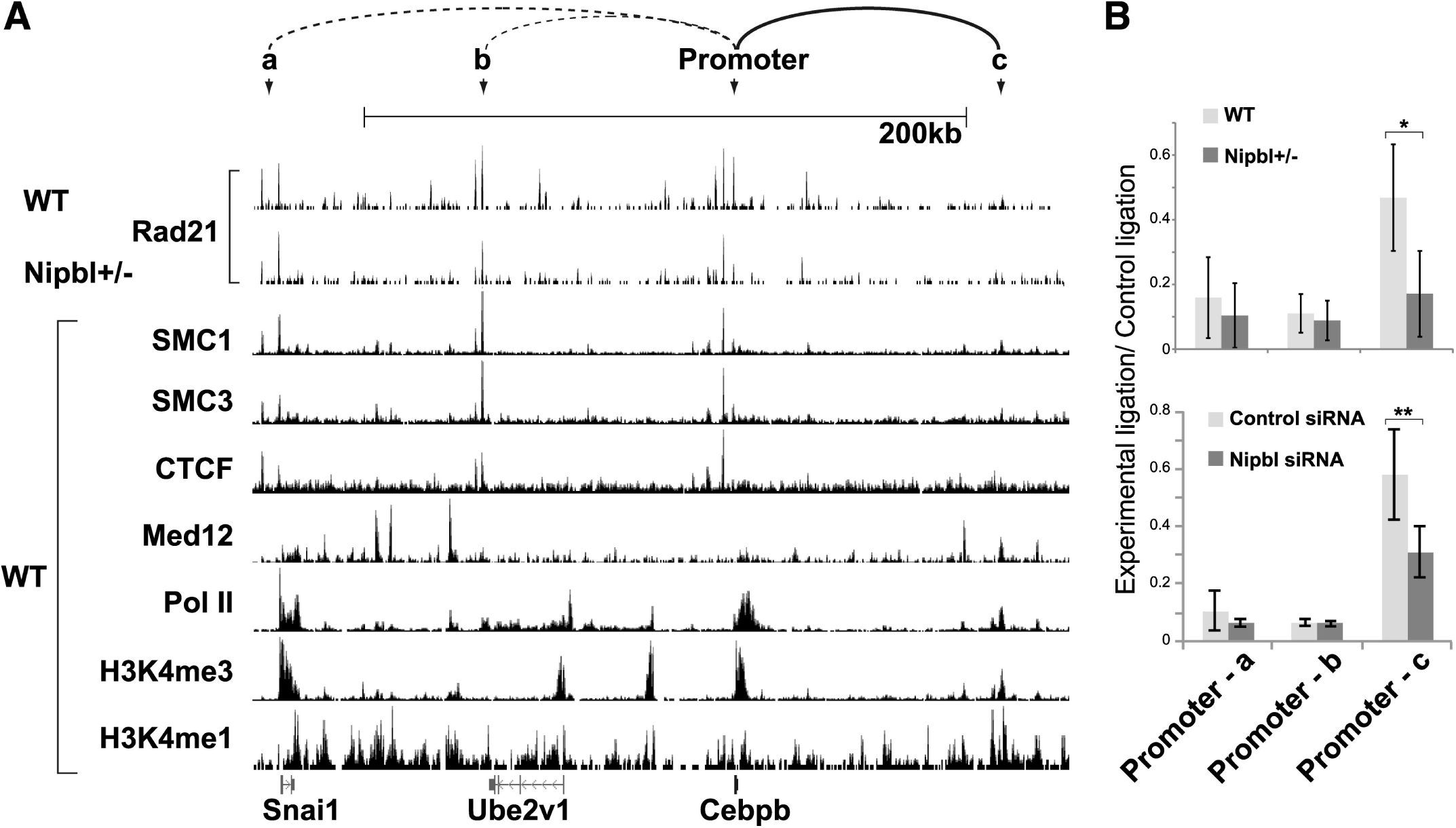
The long distance interaction involving the *Cebpβ* promoter is decreased in *Nipbl* +/- MEFs. **A.** Comparison of Rad21 binding peaks in wild type (WT) and *Nipbl* +/- mutant MEFs with SMC1 and SMC3, CTCF, and Mediator subunit 12 (Med12) [40] (GSE22562), pol II (GSE22302), H3K4me3 (GSE26657), and H3K4me1 (GSE31039) in WT MEFs in the genomic region surrounding the *Cebpβ* gene. The positions of primers for the 3C analysis (a, b, c and the promoter as the bait) are indicated. These regions were chosen based on the overlapping peaks of cohesin and CTCF, and/or cohesin, pol II and Med12 with H3K4me1/me3. The interaction observed by 3C in (B) is shown in a solid line and other interactions examined but weak are shown in dotted lines at the top. **B.** The 3C analysis of *Cebpβ* promoter interactions with regions a, b, and c (as indicated in (A)). The chromatin interactions between WT and Nipbl mutant MEFs (top panel) and between control and Nipbl siRNA-treated MEFs (bottom) were quantified and normalized as described in EXPERIMENTAL PROCEDURES. *p-value<0.01. **p-value<0.05.

## DISCUSSION

In this study, we used MEFs derived from *Nipbl* heterozygous mutant mice to analyze the effect of *Nipbl* haploinsufficiency (the primary cause of CdLS) on cohesin binding and its relationship to gene expression. We found a genome-wide decrease in cohesin binding even at CTCF sites and repeat regions, indicating the high sensitivity of cohesin binding to even a partial reduction of the Nipbl protein. Importantly, the expression of genes bound by cohesin, particularly at the promoter regions, is preferentially altered in response to Nipbl reduction. While some genes are activated, the majority of cohesin-bound genes are repressed by decreased cohesin binding, indicating the positive role of cohesin in this context. This is consistent with the significant enrichment of H3K4me3 at the promoters of cohesin-bound genes. Our results indicate that more than 50% of genes whose expression is altered significantly in *Nipbl* haploinsufficient cells are cohesin target genes directly influenced by decreased cohesin binding at the individual gene regions. One consequence of reduced cohesin binding at the promoter region is a decrease of a specific long-distance chromatin interaction, raising the possibility that cohesin-dependent higher-order chromatin organization in the nucleus may be globally altered in CdLS patient cells.

### Nipbl functions in cohesin loading at both CTCF and non-CTCF sites

In mESCs, it was suggested that Nipbl is involved in cohesin binding to only a subset of cohesin binding sites, which are largely distinct from CTCF-bound sites [40]. However, we found that Nipbl binds to, and its haploinsufficiency decreased cohesin binding to, CTCF sites in MEFs. A similar decrease of cohesin binding was observed at both CTCF insulators and non-CTCF sites in the *β-globin* locus in *Nipbl* +/- fetal mouse liver [35]. Furthermore, during differentiation in mouse erythroleukemia cells, both Nipbl and cohesin binding is concomitantly increased at these sites [35]. Therefore, while cohesin was suggested to slide from the Scc2 (Nipbl homolog)-dependent loading sites in yeast [79, 80], Nipbl is present and appears to directly affect cohesin loading at CTCF sites in mammalian cells. Nipbl, rather than cohesin, interacts with Mediator and HP1, and appears to recruit and load cohesin onto genomic regions enriched for Mediator and HP1 for gene activation and heterochromatin assembly, respectively [40, 45]. In contrast, cohesin, and not Nipbl, primarily interacts with CTCF [45, 81]. Thus, for cohesin binding to CTCF sites, we envision that cohesin initially recruits Nipbl that in turn stably loads cohesin onto CTCF sites.

A recent study indicated that almost all CTCF sites are bound by cohesin in primary mouse liver [43]. In MEFs, however, we found that ∼42% of CTCF-bound sites appear to be cohesin-free. Furthermore, there is less overlap of cohesin and CTCF in the promoter regions compared to the intergenic regions, and little correlation between CTCF binding to the promoter and gene expression changes in *Nipbl* mutant cells was observed. Thus, in contrast to the cooperative function of cohesin and CTCF at distantly located insulator sites [36], cohesin and CTCF appear to have distinct functions at gene promoters. Distinct gene regulatory functions of CTCF and cohesin have also been reported in human cells [41]. Further study is needed to understand the recruitment specificity and functional relationship of cohesin and CTCF in gene regulation.

### How does Nipbl haploinsufficiency affect cohesin target gene expression?

One mechanism of cohesin action in gene regulation is to mediate chromatin loop formation [35, 40]. Increased Nipbl and cohesin binding correlates with the induction of the enhancer-promoter interaction and robust gene activation at the β-globin locus [35]. Depletion of cohesin resulted in decreased enhancer-promoter interactions and downregulation of globin genes [35]. Similarly, *Nipbl* haploinsufficiency results in less cohesin binding and decreased promoter-enhancer interactions and *β-globin* gene expression [35]. In the current study, we also found that the cohesin-bound promoter of one of the target genes, *Cebpβ,* is involved in a long-distance chromatin interaction with a putative enhancer, which is decreased in *Nipbl* mutant cells, consistent with the decreased gene expression. Thus, *Nipbl* haploinsufficiency affects cohesin target gene expression by decreasing cohesin-mediated chromatin interactions.

It should be noted, however, that not all genes that we examined showed significant long-distance chromatin interactions involving cohesin-bound promoters. While this may be because we did not test the correct enhancer regions, it also suggests that cohesin may promote gene activation by a mechanism(s) other than by mediating long-distance promoter interaction. One possibility is gene looping. In *S. cerevisiae*, the promoter and terminator regions of genes interact with each other, which was thought to facilitate transcription re-initiation [82]. Although cohesin is often found at the promoter and terminator regions of genes in MEFs we failed to obtain any evidence for the involvement of these sites in gene looping with our limited analysis. Thus, how (or whether) cohesin at the promoter may regulate gene transcription in a loop formation-independent manner is currently unclear.

Cohesin binding to the gene body regions is found at many of the cohesin target genes. This may represent the cohesin binding at intragenic enhancer elements or may be related to Pol II pausing [29]. While cohesin was shown to facilitate Pol II elongation in Drosophila [83-85], cohesin together with CTCF in the intragenic region was found to cause Pol II pausing at the *PUMA* gene in human cells [86], suggesting that cohesin can have both positive and negative effects on transcriptional elongation in a context-dependent manner. Furthermore, not all the cohesin-bound genes changed expression in *Nipbl*+/- MEFs, echoing this notion that the effect of cohesin binding on gene expression is context-dependent. What determines the effects of cohesin binding at individual binding sites on gene expression requires further investigation.

### The role of cohesin in the maintenance of gene expression

While there is now strong evidence for cohesin’s role in chromatin organization and gene activation, whether cohesin is involved in initiation or maintenance of gene activation is less clear. Enrichment of cohesin binding at the transcription start sites and termination sites was observed previously in mouse immune cells with no significant correlation to gene expression [30]. Our genome-wide analysis also revealed that cohesin binding to the gene regions has no obvious relationship to the level of gene expression in wild type MEFs. And yet, a decrease in cohesin binding is associated with a tendency to downregulate these genes, indicative of the positive role of cohesin on gene expression, consistent with the enriched presence of H3K4me3 in promoter regions. We speculate that cohesin may not be the primary determinant of gene activation, but rather cohesin binding may be important for maintaining gene expression status initially determined by sequence- and cell type-specific transcription factors. Similarly, enrichment of bivalent histone modifications in the promoters of cohesin-bound genes with very low expression suggests that cohesin also contributes to the maintenance of the poised state of these genes.

### Nipbl haploinsufficiency vs. cohesin mutation

There are two different cohesin complexes in mammalian somatic cells that differ by one non-SMC subunit (i.e., SA1 (STAG1) or SA2 (STAG2)) [87, 88]. A recent report on SA1 knockout mice revealed some phenotypic similarity to what is seen in mice with *Nipbl* haploinsufficiency [67]. Interestingly, the *SA1* gene is one of the cohesin target genes that is slightly upregulated in *Nipbl* mutant cells [19]. Thus, together with the compensatory increase of *Nipbl* expression from the intact allele, there appears to be a feedback mechanism that attempts to balance the expression of *Nipbl* and cohesin in response to *Nipbl* mutation. The fact that upregulation was observed with the *SA1*, but not *SA2*, gene may reflect the unique transcriptional role of SA1 [67]. Interestingly, however, only 10% of genes altered in *Nipbl* mutant MEFs are changed significantly in *SA1* KO MEFs [67]. This discrepancy may, as observed in Drosophila [89], reflect the different effects of decreased binding versus complete knockout of a cohesin subunit on target gene expression. It could also be a result of the decreased binding of the second cohesin complex, cohesin-SA2.

Cohesin binding was relatively uniformly decreased genome-wide in *Nipbl* haploinsufficient cells with no significant redistribution of cohesin binding sites. Point mutations of different subunits of cohesin cause CdLS and CdLS-like disorders with both overlapping and distinct phenotypes compared to CdLS cases caused by NIPBL mutations [9, 10, 13]. Non-overlapping effects of downregulation of different cohesin subunits have been reported in zebrafish [20, 26]. This may reflect an unequal role of each cohesin subunit in gene regulation and it is possible that some of the cohesin target genes may be particularly sensitive to a specific cohesin subunit mutation. For example, similar to the TBP-associating factors (TAFs) in TFIID [90], cohesin subunits may provide different interaction surfaces for distinct transcription factors, which would dictate their differential recruitment and/or transcriptional activities. Furthermore, recent studies provide evidence for cohesin-independent roles of NIPBL in chromatin compaction and gene regulation [27, 28, 91]. Thus, disturbance of cohesin functions as well as impairment of cohesin-independent roles of NIPBL may collectively contribute to CdLS caused by NIPBL mutations.

## CONCLUSIONS

Our results demonstrate that cohesin binding to chromatin is highly sensitive genome-wide (both at unique and repeat regions) to partial Nipbl reduction, resulting in a general decrease in cohesin binding even at strong CTCF sites. Many genes whose expression is changed by Nipbl reduction are actual cohesin target genes. Our results suggest that decreased cohesin binding due to partial reduction of NIPBL at the gene regions directly contributes to disorder-specific gene expression changes and the CdLS phenotype. This work provides important insight into the function of cohesin in gene regulation with direct implications for the mechanism underlying *NIPBL* haploinsufficiency-induced CdLS pathogenesis.

## ACKNOWLEDGEMENT

We thank Dr. Alex Ball for critical reading of the manuscript.

## ABBREVIATIONS

CTCF: CCCCTC-binding factor
3C: Chromatin conformation capture (3C)
ChIP: Chromatin immunoprecipitation
CdLS: Cornelia de Lange Syndrome
FDR: false discovery rate
H3K4me3: histone H3 lysine 4 trimethylation
H3K4me1: histone H3 lysine 4 monomethylation
H3K27me3: histone H3 lysine 27 trimethylation
KS test: Kolmogorov-Smirnov test
MEFs: mouse embryonic fibroblasts
mESCs: mouse embryonic stem cells
Nipbl: Nipped-B-like
Pol II: RNA polymerase II
RPKM: reads per kb per million total reads
siRNA: small interfering RNA
TSS: transcription start site
TTS: transcription termination site.

